# Neoantigens and Stochastic Fluctuations Regulate T Cell Proliferation in Primary and Metastatic Malignant Brain Tumors

**DOI:** 10.1101/2025.04.23.650340

**Authors:** Maheshwor Poudel, William C. Stewart, Ciriyam Jayaprakash, Jayajit Das

## Abstract

Brain cancer is one of the most aggressive forms of cancer in the central nervous system occurring as primary or metastatic tumors. Sequencing of resected tissues from glioblastoma (GBM) and brain metastases (BrMET) reveals high heterogeneity in neoantigens and T cell receptor (TCR) repertoires. Our analysis of published sequencing data in different spatial regions of tumors GBM and BrMET patients show the presence of T cell clones of sizes with a heavy right-tailed distribution spanning several orders of magnitude (e.g., 1 – 1000 cells) with a few (<10) large clone sizes and many small clones. We investigated how neoantigens in the tumor microenvironment (TME) drive T cell expansion in GBM and BrMET by developing a mechanistic mathematical model based on the interaction of T cells and the neoantigens that incorporates their stochastic proliferation in the immunosuppressive environment and trained it to predict the emergence of T cell clones in different spatial regions. The model accurately predicts the distribution of observed T cell clone sizes and reveals that the strength of interaction between TCR and neoantigen-MHC complex and stochastic T cell proliferation crucially regulates T cell expansion in the TME. It also suggests higher rate of T cell proliferation BrMET compared to GBM. An extended version of the model predicts the ability of individual neoantigens to generate T cell clones in the periphery in patients receiving personalized neoantigen vaccines. Our model may facilitate the discovery of improved peptide combinations in neoantigen vaccine studies.

**Significance Statement:** Neoantigen-driven T cell responses are key to immune defense against solid tumors. Multi-region sequencing of brain tumors reveals spatial heterogeneity in neoantigens and T cell repertoires. To understand whether neoantigen-driven T cell expansion underlies the TCR repertoire heterogeneities, we developed a stochastic, mechanistic model of T cell proliferation using published TCR and neoantigen data from primary and metastatic brain tumors. The model accurately predicts clone size distributions, showing faster T cell proliferation in metastases and stronger responses to clonal (shared) neoantigens than to private (region-specific) ones. The model is extended to describe T cell clonal expansion in the periphery in response to neoantigen vaccine in glioblastoma patients. This framework may help design optimal peptide combinations in neoantigen vaccine development.

## Introduction

Brain cancer is a difficult to treat cancer and patients seldom survive beyond two years(1). Tumors in the brain can originate in the central nervous system or arise from migrating tumor cells from carcinomas of other sites in the body(2). Glioblastoma (GBM) is the most common type of malignant primary brain tumor(3), and brain metastasis (BrMET) can develop from many cancers including melanoma, and colorectal cancers(4). The brain tumor microenvironment (TME) is marked by its immune suppressive nature(5, 6) and poor infiltration of lymphocytes.

Studies using genome sequencing(7, 8), single cell proteomic(9) and transcriptomic(10, 11) technologies across multiple regions of tumors have revealed extensive intra-tumoral heterogeneity in the genetic make-up, such as non-synonymous mutations, in patients with GBM and BrMET. Intra-tumoral variations of neoantigen expressions(7) can enable tumor cells to evade natural immuno-surveillance (12), and contribute to reduced patient responsiveness to immunotherapy(12, 13). Whole exome sequencing (WES) and single cell sequencing of RNA and T cell receptors (TCRs) in multiple tumor regions in GBM and BrMET patients by Schaettler et al. (7) revealed increased intra-tumoral heterogeneity at the genomic and neoantigen levels in GBM than BrMET. Neoantigens in BrMET patients contained higher proportions of “clonal” (i.e., shared across all intra-tumor regions) neoantigens whereas higher proportions of “subclonal” (i.e., shared across a subset of intra-tumor regions) and “private” (i.e., present only in a specific tumor region) (7) were found in GBM patients. The TCR sequencing data reported by Schaettler et al. (7) showed a wide range of T cell expansion across different tumor regions in BrMET and GBM patients.

T cells are key orchestrators of the adaptive immune system which mount response to specific antigens derived from tumors or pathogens(14). The T cells in the periphery of the body are enormously diverse (∼10^8^ different T cells with unique TCRβ sequences(15, 16)) where individual T cells express unique T cell receptors (TCRs). TCRs can bind with specific peptide (p) fragments derived from pathogens or tumors when those are bound to MHC (class I and II) molecules (which we write here as pMHC complex) expressed on antigen presenting cells (APCs). A strong interaction between TCR and pMHC expressed on APCs can drive T cell proliferation(14), and neoantigen specific T cell effector functions play a major role in anti-tumor immune response and shaping tumor immunoediting(17). In non-small-cell lung cancer (NSCLC), statistical analysis of multi-region genomic and TCR sequencing data demonstrated neoantigen specific expansion of T cells in the TME(18). However, other studies in NSCLC and colon cancer showed the presence of tumor infiltrated T cells that were not specific to neoantigens but recognized many unrelated epitopes from viral infections such as EBV or CMV(19). These T cells also known as bystander T cells are speculated to be drawn into the TME by the presence of inflammation.

To this end, by combining a mechanistic and stochastic model of T cell clonal expansion with analysis of genomic, transcriptomic, and TCR sequencing data reported by Schaettler et al. (7), we investigated whether T cell clones observed in different regions of the brain tumor arise due to the antigen-driven proliferation of neoantigen specific T cells. A quantitative mechanism predicting the origin of T cell clones will be valuable for the improvement of neoantigen based therapies against brain tumors. Our modeling framework is inspired by previously reported analysis of T cell repertoire and mechanistic modeling of T clonal expansion in infection and homeostasis(20–24). By analyzing TCR sequences in different tumor regions in BrMET and GBM patients we find heavy tailed distributions of the sizes of T cell clones where the average clone size is larger in BrMET than in GBM patients. Our mechanistic model, which combines interactions between neoantigens and TCRs, the immunosuppressive nature of the tumor microenvironment in brain tumors, and the stochastic dynamics of T cell proliferation, accurately predicts the observed heavy-tailed distributions of T cell clone sizes. We find that the rate of T cell proliferation is higher in BrMET patients compared to those with GBM, and that clonal neoantigens, as opposed to private or subclonal ones, drive T cell proliferation at a higher rate.

Finally, by integrating a modified version of our mechanistic model for the TME with data from a recent clinical trial of a neoantigen vaccine in GBM patients (23), we quantified the contribution of individual neoantigens to T cell clonal expansion in the periphery of vaccinated patients. The framework can be applied to analyze and predict neoantigen driven T cell clonal expansion in other solid cancers.

## Results

### The size of T cell clones in the TME in different spatial regions in GBM and BrMET follow distributions with heavy tails

A T cell expresses about 50,000 TCR molecules on the cell surface(25) where the TCR is composed of α and β chains with unique sequences of amino-acid residues(14). The number of T cells expressing the same TCR, i.e., the size of a T cell clone can contain information regarding the proliferation of T cells in response to antigen stimulation. To estimate the size of T cell clones, we computed the frequency of occurrences of sequences of the CDR3 region of the TCRβ chain obtained from different spatial regions of resected tumor samples in BrMET or GBM patients reported by Schaettler et al.(7). Three or four distinct intra-tumor regions in a resected tumor tissue were profiled by Schaettler et al.(7) in 15 GBM and 11 BrMET patients using whole exome-, RNA-, and TCR-sequencing. Assuming any T cell, roughly expresses the same number of TCR molecules/cell, the copy number *M_i_* of a unique CDR3β chain sequence, indexed by *i* (=1, 2, .., N), is proportional to the number of T cells that belong to the clone. We assumed the proportionality constant to be unity for simplicity. The index *i* will be used interchangeably to denote a TCR containing a CDR3β chain sequence #*i* or to denote the T cell expressing the TCR#*i*. Following this notation, the size of the clone formed by T cell#*i* is given by *M_i_*. We computed the frequency of TCR clones of size *l* (or *f_l_*) in a spatial region in the TME of an individual patient by counting the number of size *l* clones that are present in all the measured TCR sequences present in a spatial region, i.e., 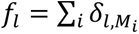. We normalize *f_l_* such that ∑*_l_f_l_* = 1. The clone sizes varied across different spatial regions in individual patients and across patients (**Figure 1A**). The average clone size in BrMET patients (=2.3) is larger than that in the GBM patients (=1.8) (**Figure 1A**), and the clone sizes show a wider variation in BrMET patients compared to the GBM patients (**Figure 1B**). Next, we analyzed the clone size distributions *f_l_* in different spatial regions in BrMET and GBM patients. The variation of *f_l_* with *l* showed the following trends: (1) *f_l_* varied with *l* as a power law (*f_l_* ∝ *1/l^α^* ) with an exponent of α ∼ 1.2-4.3 for over two decades for many spatial regions across the GBM and BrMET patients (**Figure 1C**). The power law distribution here reveals the presence of a wide range of clone sizes, where typically just a few large clone sizes (>100 cells) will co-exist with several hundred smaller clone sizes. Power law distributions have been observed in many areas such as financial net worth of individuals and magnitudes of earthquakes and may point to novel mechanisms underlying the distribution in certain situations or impose specific constraints on models explaining the behavior(26). (2) We did not find any statistically significant difference in the means of the exponent *α* computed from different regions in the same patient or across BrMET and GBM patients (**Figure 1D**). The range of *l* for each power law behavior could be less than a decade for several regions and patients, thus the data might not represent true power law decays which span several decades of *l*. Overall, the analysis of the T cell clone sizes showed a higher average value in BrMET compared to GBM patients, and the distribution of the clone sizes showed heavy tails indicating the co-existence of a wide range of T cell clone sizes in most of the spatial regions.

**Figure 1.**
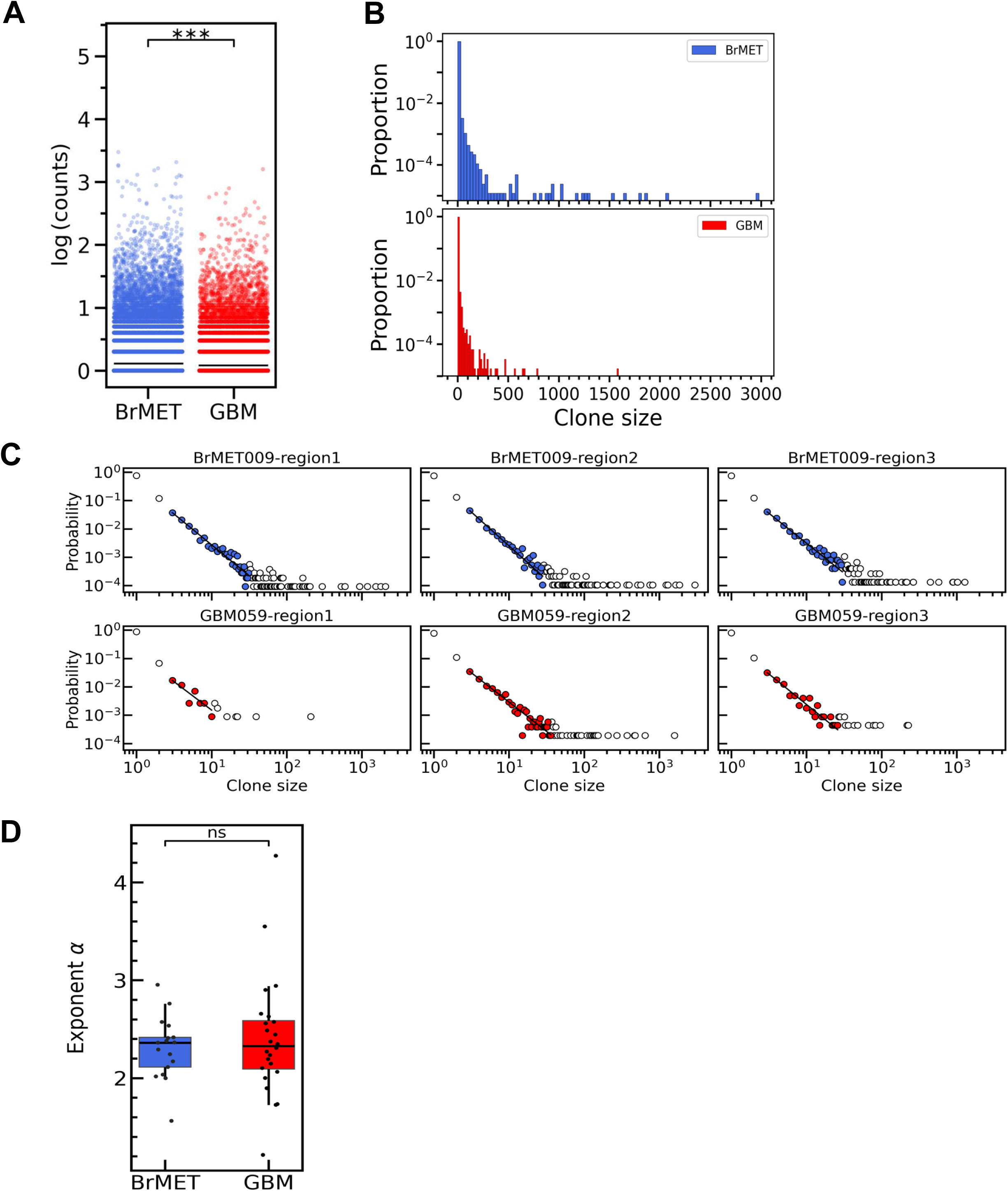
T cell clone sizes show broad heavy tailed distributions in BrMET and GBM patients. **(A)** Shows T cell clone sizes in log scale estimated in spatial regions in BrMET and GBM patients. The mean clone size in BrMET is significantly higher than that in GBM. Significance is determined by two-sided Welch’s t-test. **(B)** Shows the normalized probability distribution of T cell clone sizes shown in (A). The clone sizes in BrMET show a wider variation and an increased frequency for the presence of clone sizes over 100 compared to the GBM counterparts. (**C**) Shows power law (∝ 1/*l^α^*) fits to the clone size distributions in three different spatial regions of a BrMET and a GBM patient. The data (shown in color) that fit the power law span clone sizes over a decade. (**D)** The box plots of the power law exponent (*α*) estimated by fitting the clone size distributions in different spatial regions in BrMET and GBM patients. The mean value of α does not show any statistically significant different between the BrMET and the GBM patients. Significance determined by two-sided Welch’s t-test.

### Development of a stochastic and mechanistic model to describe TCR clone size distribution

Proliferation of a T cell requires stimulation of TCRs by neoantigen bound MHC molecules expressed on antigen presenting cells (APCs) such as dendritic cells. Schaettler et al. predicted about 935 (BrMET) and 448 (GBM) neoantigen candidates using the software *pVACtools* (27) and WES data in each spatial region of the tumor tissue samples resected from BrMET and GBM patients. Chain and colleagues(18) showed for neoantigen-driven T cell expansion in non-small cell lung cancer (NSCLC) the number of distinct T cell clones is positively correlated with the number non-synonymous mutations in a tumor. Since the number of distinct neoantigens is usually proportional to the number of non-synonymous mutations in a tumor region(28), we computed the correlation between the number of distinct T cell clones and the number of distinct neoantigens which showed negligible correlation (**Figure S1A**). The lack of this correlation could be due to the differences in the strength of interactions between neoantigens and TCRs in different regions (**Figure S1B**). We reasoned if the observed T cell clones result from proliferation of T cells in the TME due to TCR interacting with neoantigen-MHC molecules, the TCR clone sizes should be correlated with the total effective strength of the interactions. We created a variable for describing the TCR-neoantigen-MHC interactions where a T cell expressing TCR #*i* interacts with an APC expressing neoantigen *u* with a strength *s_iu_*. We assumed any T cell (e.g., T cell#*i*) in a spatial region interacts with all the neoantigens in the same spatial region with equal probability, thus the propensity (or the rate) of proliferation for T cell #*i* is proportional to the net strength of interaction experienced by T cell #*i*, *r*_*i*_ = ∑_*u*_ 𝑠_*iu*_(**Figure 2A**). We estimated *s_iu_* by using the TCRβ and neoantigen sequences in bioinformatic tools *PanPep*(29) and *ERGO-II*(30). The bioinformatic tools (*PanPep* and *ERGO-II* input the CDR3β and the neoantigen sequences to generate an estimated value for *s_iu_. ERGO-II* uses two types of training where the tool is trained on *VDJdb* or *McPAS* datasets. The magnitudes of *r_i_* computed for different spatial regions across patients show region to region variations in individual patients and patient-to-patient variations (**Figure 2B and Figures S3A-B**). The means of {*r_i_*} in different spatial regions in individual BrMET and GBM patients do not show appreciable difference (**Figure 2B**). However, the means of {*r_i_*} collected from the spatial regions and patients show a higher magnitude in BrMET patients compared to GBM patients (**Figure 2C**) suggesting the presence of higher affinity neoantigens in BrMET compared to GBM patients. However, when we computed the correlation between the estimated *r_i_* and the clone size 𝑀_*i*_ for the TCR clones, they showed poor correlation (**Figure 2D and Figures S3C-D**). We reasoned that the lack of correlation could arise from several potential sources. First, the TME in brain tumor usually contains few professional APCs such as dendritic cells (31, 6) which can efficiently present neoantigens to T cells and lead to proliferation. Second, the brain TME is immunosuppressive where the immune suppression is mediated by a variety of factors including macrophages(5, 6), cytokines such as IL-10(32) and TGFβ which could inhibit T cell activation leading to proliferation. In this situation, a T cell needs to search or wait for certain time τ before it receives a sufficient level of stimulation by neoantigen-MHC leading to its proliferation. Third, the proliferation of a T cell upon interacting with a pMHC is intrinsically stochastic(33, 34), i.e., the same TCR-pMHC interaction may not always result in the same outcome (proliferation or no proliferation). This randomness describes the inherent uncertainties in cell decision processes.

**Figure 2.**
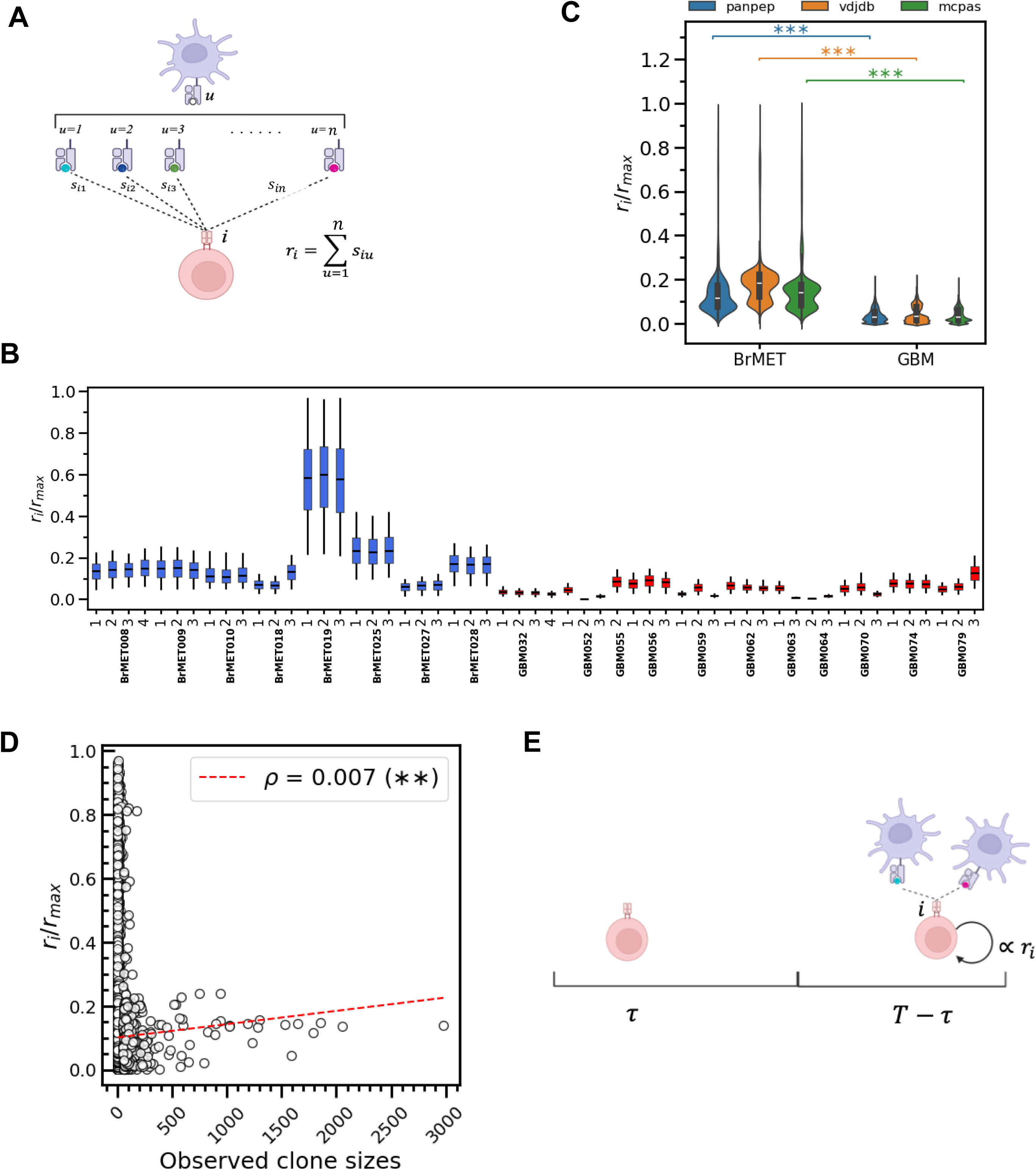
Development of a stochastic and mechanistic model to describe TCR clone size distribution. **(A)** Schematic depiction of the calculation of propensity to proliferate for an 𝑖^*th*^ TCR. The 𝑖^*th*^TCR is assumed to interact with all possible neoantigens {𝑢_*i*_, 𝑖 = 1, 2, … , 𝑁} such that the proliferation strength of the respective T cell is aggregated sum of pairwise (*iu*) interactions i.e., *r_i_* = ∑*_u_i__* 𝑠*_iu_* **(B)** Box plot of propensity to proliferate *r*_*i*_ (scaled by a common factor and calculated using *PanPep*) for all BrMET patients (in blue) and GBM patients (in red). For a patient within the regions analyzed, the *r*_*i*_ values do not show any appreciable differences. **C,** Box plots of the propensity to proliferate between BrMET and GBM groups calculated from *PanPep* and *ERGO-II* (trained on *VDJdb* or *McPAS* databases). Note {*r*_*i*_} values were divided by the corresponding *r*_561_ (=max{r_i_}_all BrMET+GBM patients_) values for *PanPep* and *ERGO-II* (*VDJdb* / *McPAS*) estimations. BrMET group has higher propensity to proliferate than GBM such that possibly BrMET has higher affinity T cells than GBM. Significance determined by two-sided Welch’s t-test. (**D)** Scatter plot between propensity to proliferate for every TCRs versus their observed clone sizes. Regression line shown in red. Significance determined from Pearson R correlation test. Although, the test signifies that the correlation between these two quantities is significant, however the correlation is negligibly weak. (**E)** Model of T cell activation and proliferation in the brain tumor TME. We assume that a T cell enters the TME and resides for time 𝑇. The T cell waits for an interval 𝜏 ≤ 𝑇 to search for pMHC complex and interact with it for activation. Upon activation, the T cell clone expands during the interval 𝑇 − 𝜏.

Both the above factors as well as other sources such as inaccuracies in the prediction of the strength TCR-pMHC interaction by bioinformatic tools(16, 35) can mask any direct relation between the strength of the TCR-neoantigen-MHC interaction and the TCR clone size. However, the waiting of the T cell before it can proliferate and the stochastic nature of the proliferation can be accounted for in a simple mechanistic model (**Figure 2E**), so we investigated if including these effects in a model can lead to excellent prediction the TCR clone size distributions in the TME. In our model, we assume T cells enter the TME and spent a total duration of time *T* in any spatial region until the tumor was resected. A T cell waits a time interval *τ* to receive stimulatory signals in the TME in the model, and then proliferates in the duration *T-τ*. We consider the waiting time *τ* to be distributed in an exponentially increasing function with time in the inhibitory region, 𝑝(𝜏) ∝ exp(𝑘𝜏), i.e., the longer the T cell waits the more likely it becomes for it to experience a productive interaction to undergo proliferation. The T cell proliferation is modeled using a Markov process, and the probability 𝑝(𝑀_*i*_, 𝑇; 𝑥_*w*_, 𝑥_*p*_*r*_*i*_) that a T cell#*i* will form a clone size of *M_i_* in the time interval *T* can be calculated analytically and exactly (details in Materials and Methods section). The model parameters *x_w_* and *x_p_* are estimated by using the *r_i_* values calculated from *PanPep* or *ERGO-II*, and the measured T cell clone size data in GBM and BrMET patients. We describe the parameter estimation and determine potential underlying mechanisms in the next section.

### Distribution of TCR-neoantigen:MHC interactions and stochastic fluctuations describe T cell clone size distributions

We used the mechanistic model developed above to describe the size distributions of T cell clones observed in different spatial regions in individual patients. Given the counts of the T cells #*i* (or {*M_i_*}) and the strength of TCR-neoantigen-MHC interactions {*s_iu_*} for the neoantigens {*u*}found in a spatial region, we maximized the negative log-likelihood function (−ln ℒ) to estimate the parameters {*x_p_*, *x_w_*} (details in the Materials and Methods section). The proliferation rate is a product of the parameter *x_p_* and *r_i_*, the average antigen interaction strength experienced by T cell#*i*. We found that the log-likelihood function does not change appreciably as *x_w_* is varied over a wide range, thus we fixed value of *x_w_* at 100, and then estimated *x_p_* in all our calculations (details in Materials and Methods and **Figure S2**). The estimated values of *x_p_* showed lower mean value across spatial regions and patients in BrMET than GBM patients (**Figure 3A**), however, the mean value of {*x_p_r_i_*} (=𝑥_*p*_*r̅*) or the mean propensity (or the rate) of T cell proliferation across spatial regions and patients is higher in BrMET than the GBM patients implying increased T cell proliferation rate in BrMET patients (**Figure 3B**). Since macrophages present in the TME can induce inhibit T cell activation(5, 6) we checked whether the correlation of the T cell proliferation with the proportion of macrophages present in a spatial region.

**Figure 3.**
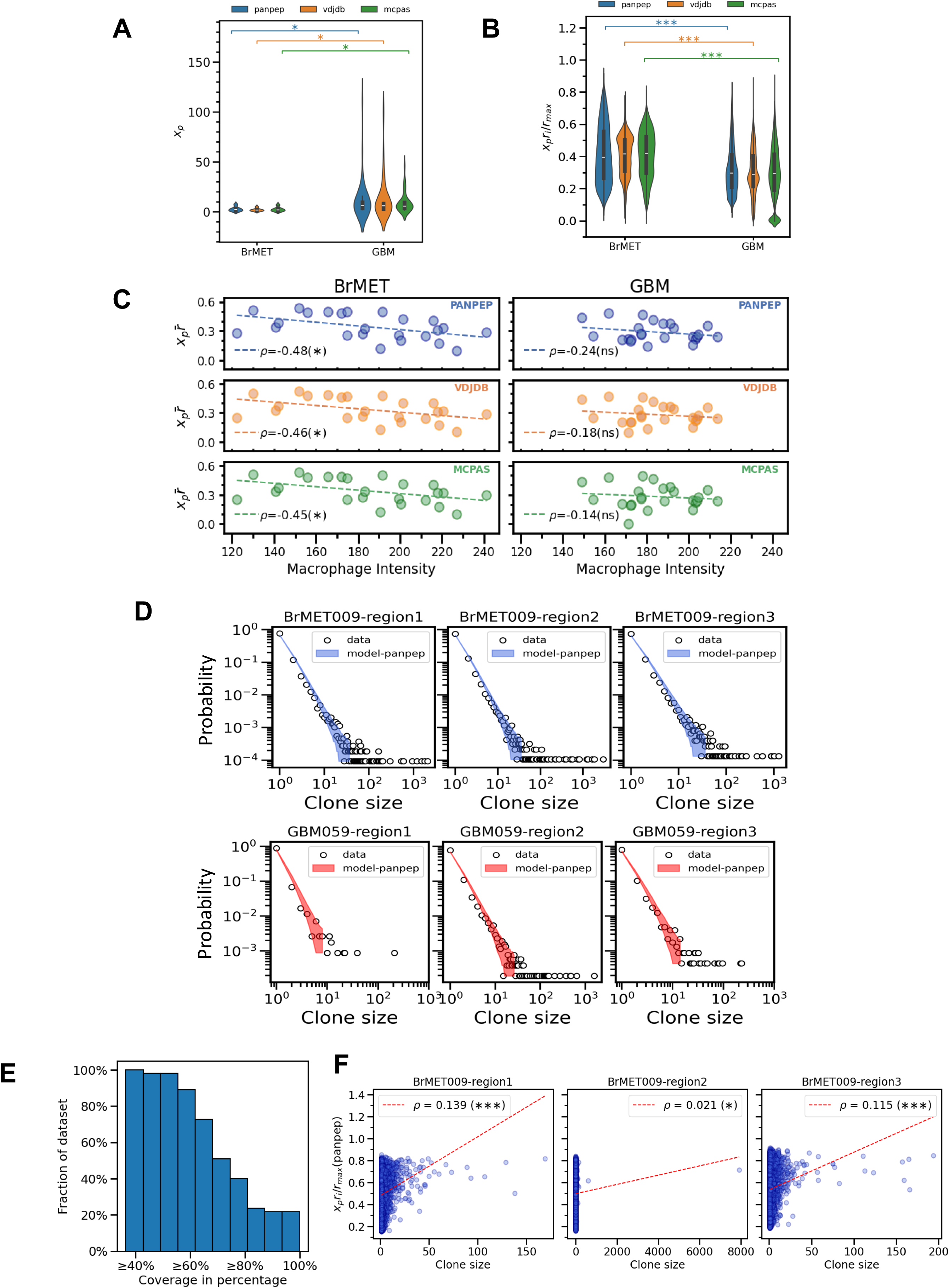
Distribution of TCR-pMHC interactions and stochastic fluctuations describe T cell clone size distributions. **(A)** Violin plot of proliferation parameter 𝑥_*p*_ between BrMET and GBM groups. The *x_p_* values showed lower mean value across spatial regions and patients in BrMET than GBM groups. Significance is determined by two-sided Welch’s t-test. **(B)** Violin plot of effective proliferation rate 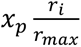 between BrMET and GBM groups. The mean propensity 𝑥_*p*_*r̅* is higher in BrMET than in GBM. The significance is qualitatively true irrespective of the bioinformatics tools we used to estimate the proliferation rate. Significance determined by two-sided Welch’s t-test. (**C)** Scatter plots between effective proliferation rate versus macrophage intensity for BrMET and GBM groups. The average macrophage intensity and the average propensity 𝑥_*p*_*r̅* across the spatial regions is negatively correlated in the BrMETs suggesting macrophages potentially lead to suppression of T cell proliferation in BrMET but not in GBMs. Significance determined from Pearson R correlation test. (**D)** Model prediction of the clone sizes for BrMET (represented here by BrMET009 patient) and GBM group (represented here by GBM059 patient). The band of prediction is obtained from 1000 samples. (**E)** Summarizes the coverage probability of model prediction for the T cell clone size data for spatial regions across patients. (**F)** Scatter plot of the effective proliferation rate versus the model predicted clone sizes (single sample/realization of the T cell clones) for BrMET009-region1. The correlation between these two quantities is negligibly weak. Significance determined from Pearson R correlation test.

Schaettler et al.(7) measured single cell RNA-seq in the spatial regions, and the number of macrophages in the spatial regions was quantified by a macrophage intensity obtained by deconvoluting the scRNA-seq data. We found negative correlation (𝜌 = −0.48, 𝑝 = 0.0018) between the average macrophage intensity and the mean propensity (𝑥_*p*_*r̅*) across the spatial regions in individual patients in the BrMET group regardless of the software used (e.g., *PanPep* or *ERGO-II*), whereas, this correlation was not significant (𝜌 = −0.24, 𝑝 = 0.25) in the GBM group of patients (**Figure 3C top panels**) suggesting macrophages potentially leads to suppression of T cell proliferation in BrMET but not in GBM.

Next, we used the joint distribution of the TCR clones (Eq. 2) in the model with estimated model parameter *x_p_* to generate many realizations of TCR clones arising due to stochastic fluctuations and then evaluated the TCR clone size distribution in each spatial region for individual patients. We compared the *f_l_* obtained from the TCR clone sizes for the measured TCR sequences in patients and configurations of TCR counts predicted by the model for the same patient. The model predictions showed excellent agreements with predictions producing coverage probability of over 70% for 51% of the patients (region by region) (**Figures 3D**, **3E**, **and Figure S4**, and more details in the Materials and Methods section). We also checked if the clone sizes of a single realization of the TCR clones generated in our model correlated with the strengths ({𝑠_*iu*_}) of the neoantigens in a spatial region in an individual BrMET or GBM patient. This single realization would be a representation of the TCR sequences observed in a spatial region in a patient, and we found, similar to that observed for the clones observed in patients (**Figure 2C**), the clone sizes showed almost no correlation with the strength of the TCR-neoantigen-MHC interactions (**Figure 3F and Figure S5**) suggesting the effect of intrinsic stochastic fluctuations in generating low or no correlations. The results point to several mechanistic features of T cell kinetics in the TME in brain tumors: (i) The T cells proliferate more in the TME of BrMET than GBM patients. (ii) The presence of macrophages in the spatial regions can inhibit T cell proliferation in BrMET patients. (iii) T cell proliferation in the TME is affected by intrinsic stochastic fluctuations in T cell replication process.

### Different neoantigens give rise to different T cell clone size distributions in the TME

Different neoantigens ({*u*}) interact with the same TCR#*i* with different strengths ({*s_iu_*}), and as we showed above the relation between the clone size of a TCR (e.g., #i) and the interaction strengths ({*s_iu_*}) of that TCR with other neoantigens is not straightforward due to the stochasticity in the T cell proliferation process and the availability of the TCR sequencing data from a single sample (**Figure 3D**). Therefore, to determine the contribution of an individual neoantigen in T cell expansion based on its interaction with the TCRs in a spatial region we developed the following scheme using our mechanistic model. We constructed a set of alternate scenarios for any spatial region where in each which we replaced the mixture of neoantigens found in that spatial region by one of the neoantigens present in the mixture. Thus, when *n* number of different neoantigens indexed by *u* (=1,..,*n*) are present in a spatial region, we constructed *n* different alternate scenarios indexed by {𝑝_1_, 𝑝_2_, . . , 𝑝_*n*_}, where in any alternate scenario denoted by 𝑝_*u*_, *n* copies of a single neoantigen denoted by *u* are present (**Figure 4A**). Next, we computed the clone size distributions for the T cells in a spatial region in each of the alternate scenarios using the model trained on data in that spatial region. The mean T cell clone size, averaged over many realizations of the T cell clones generated by the model in a tumor region, in each of these alternate scenarios showed strong positive correlation with the mean propensity (=𝑥_*p*_*r̅*) in BrMET and GBM patients (**Figure 4B and Figures S6A-B**). A larger value of the mean clone size in an alternate scenario expressing a particular neoantigen suggests a greater contribution of that neoantigen in the neoantigen repertoire in giving rise to T cell clones in a tumor region. This allowed us to rank order individual neoantigens in a patient based on their ability to give rise to increasing sizes of T cell clones in the spatial regions of the TME (**Figure 4B**). We found there exists select neoantigens where the mean propensity (=𝑥_*p*_*r̅*) in the alternate scenario containing that neoantigen by roughly the same to that of the neoantigen mixture found in the spatial region. These neoantigens in our model produced T cell clone size distributions close to that observed in the spatial region – thus the select neoantigen alone can potentially produce a T cell clone size distribution generated by a mixture of neoantigens (**Figure 4B and Figures S6A-B**). Next, we investigated if the clonal, sub clonal, or the private neoantigens differed in their abilities to generate T cell expansion in the TME of BrMET and GBM patients. Since the mean of the proliferation rate (𝑥_*p*_*r̅*) in the alternate scenarios presenting single neoantigens correlates with the mean T cell clone size, we computed the mean proliferation rates for the alternate scenarios each presenting a single neoantigen. We found that the mean proliferation rates for the clonal neoantigens are greater than sub clonal or private neoantigens in BrMET patients suggesting greater contribution of clonal neoantigens in driving T cell expansion (**Figure 4C)**.

**Figure 4.**
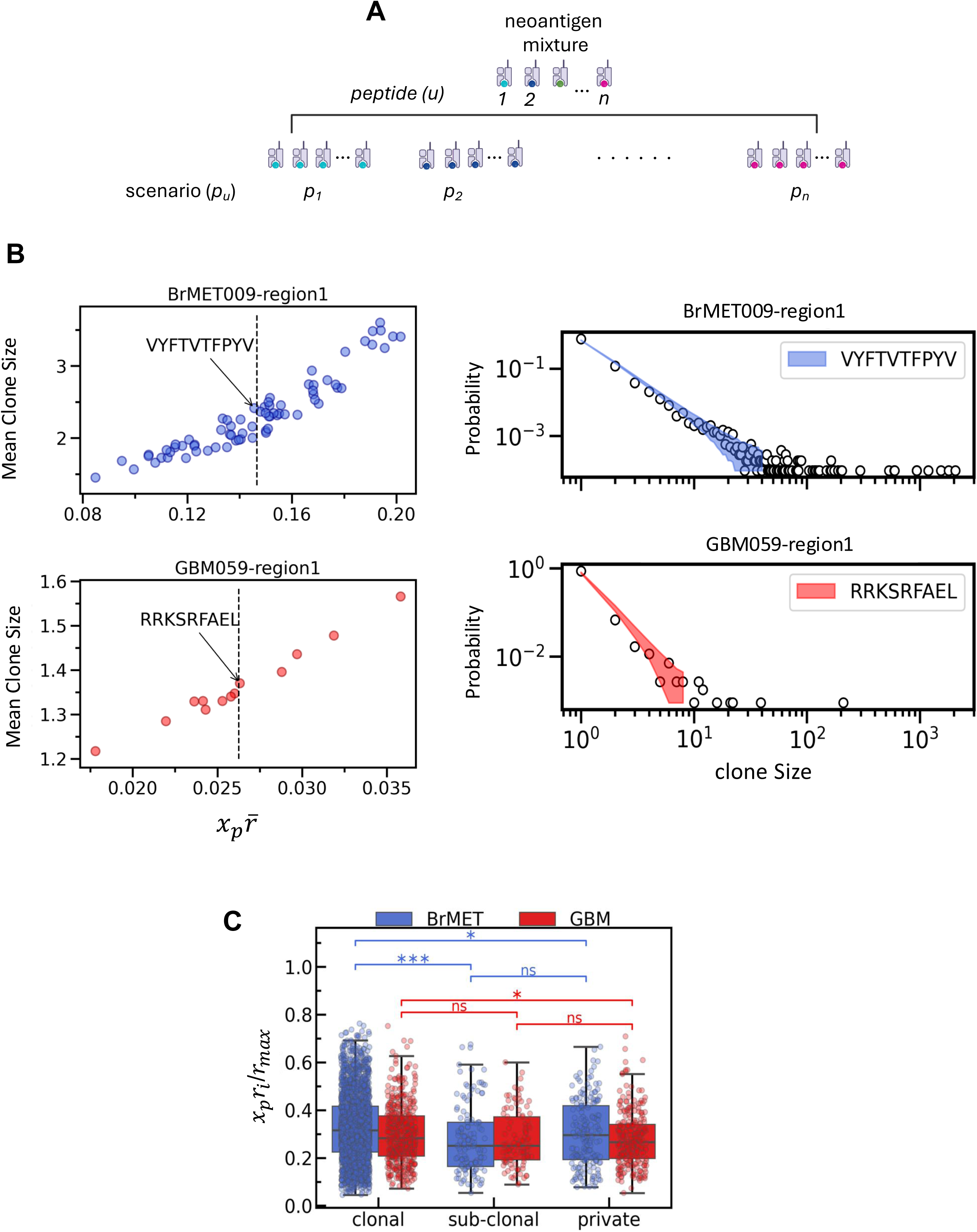
Different neoantigens give rise to different T cell clone size distributions in the TME. **(A)** Schematics of scenarios when the mixture of neoantigens that TCRs bind to is replaced by a single neoantigen one at a time. If for a spatial region there are 𝑛 neoantigens indexed as 𝑢 = 1,2, . . , 𝑛, then there are 𝑛 scenarios, each indexed as 𝑝_1_, 𝑝_2_, … , 𝑝_*n*_ of replacing the observed mixture of neoantigens by individual neoantigens 𝑢. **(B)** The mean clone size produced by each neoantigens or per scenario in A shown in left panels for patients BrMET009 (top and blue) and GBM059 (bottom and red) in region 1. The neoantigens labelled are the neoantigens which have the mean propensity closest to the mean propensity when all the peptides are considered shown in black dotted line. The right panels show the model predicted clone sizes when The TCRs are probed only by the respective neoantigens mentioned earlier. The model prediction in these scenarios is close to the model prediction in Figure 3D indicating that we can find several candidate neoantigens which alone can probe the T cell clonal expansion. **(C)** Box plots of mean propensity for every neoantigens (from *PanPep*) for BrMET and GBM groups across regions differentiated as three categories clonal, sub-clonal and private neoantigens based on their presence in the region sequenced. The clonal neoantigens show higher mean propensity than the other neoantigens irrespective of the type of tumor. Significance determined by two-sided Welch’s t-test.

### Clonal expansion of T cells in response to neoantigen vaccines in glioblastoma patients

We studied clonal expansion of T cells in the periphery in glioblastoma patients who were given neoantigen vaccines (NeoVax) as a part of a clinical trial reported by Johanns et al. (36). The personalized vaccine for an individual patient contained a mixture of neoantigens obtained from resected tumor tissues from the patient. The PBMCs were collected pre-vaccination and at post-vaccination at >60 days. We identified identical TCRβ sequences in pre- and post-vaccination samples and further analyzed the T cell clones that increased in size or remained the same size (e.g., fold change ≥1). The size distributions of these T cell clones can be fitted with a power law for a range of sizes close to a decade with an exponent about 2.3 (**Figure 5A**). The exponent does not vary appreciably between pre- and post-vaccination or across patients. The fraction of clones over size 100 increased from 53% to 57% from pre- to post-vaccination in subject#3 indicating a small increase in the expansion of large T cell clones post vaccination. We modified our mechanistic model for T cell clonal expansion in the TME to describe the clonal expansion in the periphery. Since T cells can potentially interact with the neoantigens present in the vaccine in the lymph nodes, we did not consider any delay in T cells receiving appropriate stimulation for proliferation in the model (**Figure 5B**). The T cells in the model proliferate stochastically proportional to the strength of the interaction between TCR and neoantigen-MHC. We evaluated the probability 𝑝(𝑀_*i*_, 𝑇|𝑚_*i*_, 0; 𝜆*r*_*i*_) of finding a clone of size 𝑀_*i*_ post-vaccination for a T cell#*i*, given its size 𝑚_*i*_ pre-vaccination, after a time duration *T* analytically and exactly using our model (details in Materials and Methods). The rate of proliferation *r_i_* for TCR#*i* was evaluated using *PanPep* and *ERGO-II*, and we estimated the model parameter λ using the T cell count data (details in Materials and Methods), which varied across patients(**Table S1**). We generated T cell clone size distributions at post-vaccination from the estimated probability distributions which agreed reasonably well with the data (**Figure 5B and Figures S7A-B**). Next, we quantified the contribution of individual neoantigen peptides in the vaccine mixture in generating T cell expansion using our model following a similar approach described in the previous section (**Figure 4**). Analogous to the alternate scenarios (**Figure 4A**) described in the previous section we considered alternate vaccines where we replaced the mixture of neoantigens in a neoantigen vaccine by a single neoantigen, and then used the model to predict T cell clone size distribution and the average clone size at 96, 68, and 169 days post vaccination for subject#1, subject#2 and subject#3 respectively. The rank ordering (from the largest to the lowest) of neoantigens based on the average clone sizes generated by alternate vaccines shows moderate difference between the highest and the lowest average clone sizes (**Figure 5D and Figure S7C**). The clonal neoantigens produced the largest mean T cell clone size in subjects#1-3 (note the vaccine for subject#2 contained only clonal neoantigens). Johanns et al. (36) performed ELISPOT IFNγ assay of PBMCs that were expanded in vitro by individual neoantigen peptides along with IL-2 for 12 days, and found specific clonal peptides produced the largest reactivity. The rank ordering of neoantigen peptides in giving rise to PBMC reactivity in the above assay does not agree with that shown in **Figure 5D** which could be due to a variety of factors such as differences in in vitro vs in vivo expansion, differences in IFNγ response and cell proliferation in antigen stimulated T cells(37), and inaccuracies in the bioinformatic tools *PanPep* and *ERGO-II* in estimating the strength of the interactions between TCRs and neoantigens. Overall, our modeling framework provides a systematic framework to determine relative contribution of individual neoantigen peptides.

**Figure 5.**
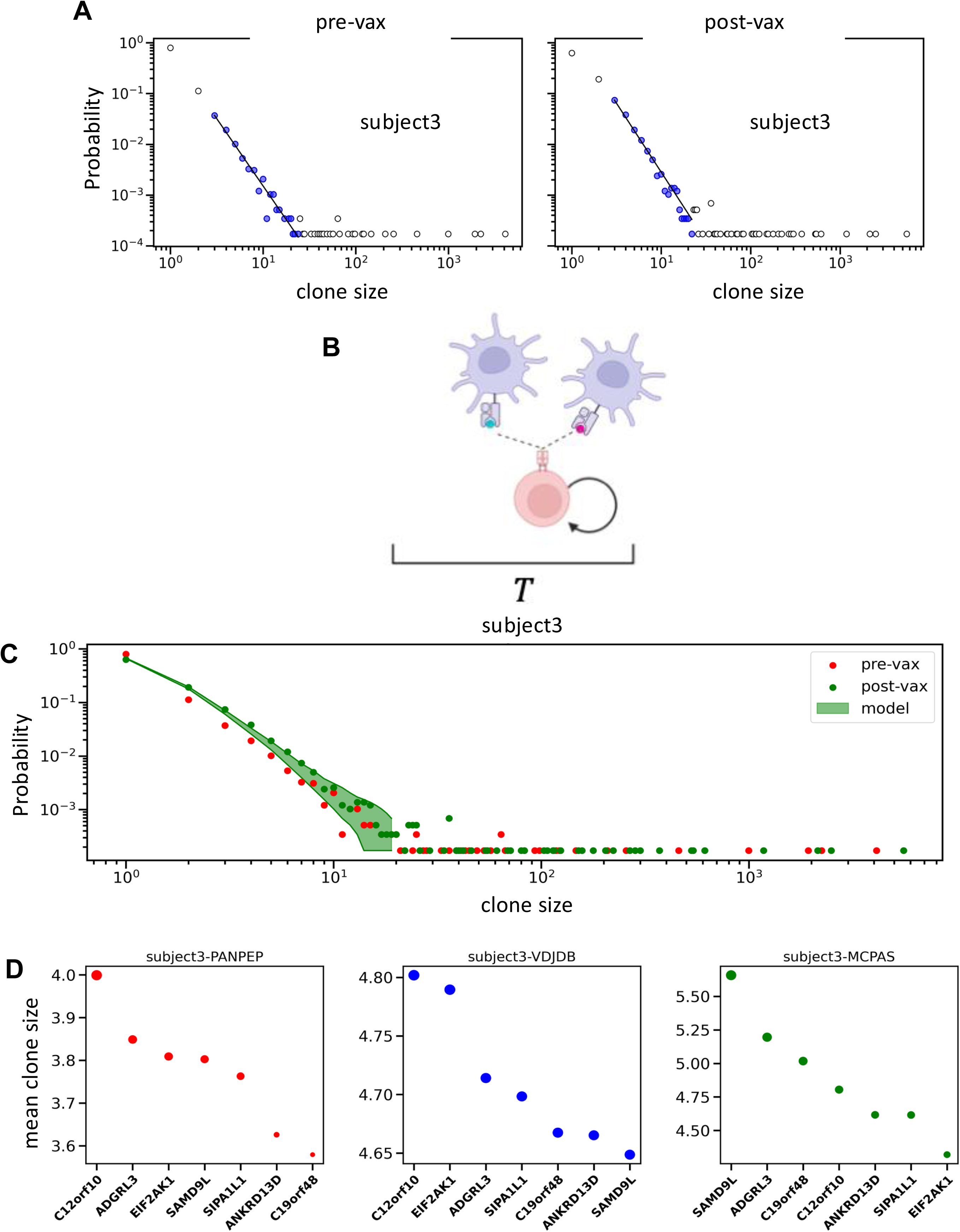
Clonal expansion of T cells in response to neoantigen vaccines in glioblastoma patients. **(A)** Power law fit (solid line) to probability distribution of observed clone sizes in subject 3 pre- and post-vaccination. The clone sizes considered in the fit are shown in blue. The exponents vary between 2.2 and 2.7. **(B)** Schematic description of the model. T cells proliferate due to stimulations received from neo-antigen presenting APCs. **(C)** Shows comparison of the model predictions for T cell clone sizes post vaccination in subject#3. (**D**) Rank ordering (ascending) of synthetic long peptides used in neoantigen vaccine in Johanns et al. based on their ability to give rise to increased T cell clonal expansion as predicted by the model. The mean T cell clone sizes are used to represent increased T cell expansion in the model (see text for details). The size of the dots represents the magnitude of mean propensity for the respective peptide. The order depends on the relative estimates of the TCR-pMHC strengths {*r*_*i*_} estimated by *PanPep*.

## Discussion

We investigated how interactions between neoantigens and the TCRs regulate antigen-driven T cell proliferation across multiple regions of primary or metastasized brain tumors. The sizes of T cell clones obtained from multiple regions in tumor tissues in GBM and BrMET patients(7) showed a heavy tailed behavior, consistent with a power law, characterized by the coexistence of numerous small (∼ 1 to 5 cells) clones alongside a few exceptionally large clones (∼100 −1000 cells). In solid cancers such as NSCLC and colon cancer, tumor infiltrating bystander T cells that are not specific to neoantigens and likely attracted by the local inflammatory response have been identified(19). Thus, the observed T cells in the brain tumor could potentially contain bystander T cells. We determined the role of antigen-specific proliferation of T cells across different regions of brain tumors underlying the emergence of observed T cell clones in GBM and BrMET patients by employing a mechanistic and stochastic model of T cell proliferation trained on TCRβ sequences and neoantigens reported by Schaettler et al. (7). We estimated the effective strength of interactions between T cells and neoantigens using two widely used bioinformatic tools *PanPep* and *ERGO-II*. However, we noticed that the size of a T cell clone is almost uncorrelated with the net interaction strength the TCRs experience with the neoantigens present in a tumor region in both BrMET or GBM patients. We reasoned that this apparent lack of correlation may arise from the intrinsic stochasticity in T cell proliferation, combined with the exponentially increasing distribution of the times T cells wait in the brain TME before receiving the necessary signals to proliferate—highlighting the immunosuppressive nature of the TME. Built on these assumptions and trained using observed T cell clone sizes in tumor regions, our mathematical model accurately predicted the heavy-tailed distributions of T cell clone sizes across various intratumoral regions in both BrMET and GBM patients. The T cell clone sizes generated from our model reproduced the negligible correlation between the size of a T cell clone and the effective strength of its interaction with neoantigens further supporting our model assumptions regarding the exponentially increasing waiting time distribution and the stochasticity in T cell proliferation. Thus, the model suggests T cell proliferation in the TME of brain tumors is driven by neoantigen-MHC and TCR interactions and is modulated by both the intrinsic stochasticity in T cell replication process and the immunosuppressive nature of the TME.

We found several key differences in the T cell response in the TME in GBM and BrMET patients. The TCR repertoire data in the patients revealed that the average T cell clone size in the tumor regions is higher in BrMET patients compared to that in GBM patients indicating an increased T cell proliferation in BrMET patients. This reasoning is supported by the larger value of the estimated average rate of T cell proliferation in our mechanistic model in BrMET patients compared to the GBM patients and suggested that neoantigen specific T cell proliferation gives rise to the observed the larger T cell clones in BrMET compared to that in GBM. Furthermore, we found increased presence of macrophages in the tumor regions correlated with lower T cell proliferation rates in BrMET patients implying inhibition of T cell activation by macrophages in the TME, in contrast, the lack of this correlation in GBM potentially indicates a qualitatively different macrophage induced suppression of T cell response in GBM. Schaettler et al. (7) found the macrophages in BrMET contained higher expression of genes associated with monocyte derived macrophages than GBM, and the macrophages in GBM were more enriched in microglia specific genes. Thus, the results suggest that the macrophages in BrMET might assume an immunosuppressive phenotype (e.g., M2) whereas in GBM the inhibition of T cells could be further modulated by factors such as cytokines such as TGFβ(5) and other unconventional modes(38). The macrophage induced inhibition of T cell activation in the TME in BrMET is consistent with the higher effectiveness of ICB therapy in metastasized brain tumors(39) compared to primary brain tumors.

We employed our model to quantify relative contribution of individual neoantigen peptides in generating T cell proliferation. In BrMET, we find that clonal neoantigens, expressed at higher proportions across multiple tumor regions, are associated with a higher rate of T cell proliferation compared to subclonal and private neoantigens. In contrast, in GBM, where subclonal and private neoantigens are expressed at higher levels than in BrMET, clonal neoantigens still drive a higher rate of T cell proliferation compared to their private counterparts.

This suggests a greater potential of using clonal neoantigens in developing personalized vaccines against GBM and BrMET. We modified our mechanistic model developed for the TME to describe T cell expansion in response to neoantigen vaccines in the periphery in GBM patients. The model showed good agreement of the clone size distributions with model predictions. The rank ordering of neoantigen peptides in their ability to generate T cell proliferation showed that clonal neoantigens produced the largest T cell proliferation in the subjects. Thus, our modeling framework can potentially be employed to design neoantigen vaccine candidates.

## Limitations

The estimation of TCR-pMHC interactions based on the sequences of CDR3β sequences of TCR, peptide, and MHC by software packages such as *PanPep* and *ERGO-II* still remains inaccurate within a wide range of ∼40-90% accuracy(35). This could produce inaccurate prediction of clonal expansion of a particular T cell expressing a unique TCRβ sequence in our model. However, the qualitative features of model results and predictions regarding T cell clone size distributions are likely to be less affected. This is further substantiated by the qualitative similarities of the results obtained using *PanPep* and *ERGO-II*. The rank ordering of T cell expansion produced by specific peptides depended on the bioinformatic tool (*PanPep* vs *ERGO-II*) and the databased they were trained on (*VDJdb* or *McPAS* datasets for *ERGO-II*) arising due to the lack of consensus in predicting the strength between the TCR and neoantigen-MHC due to the differences in underlying algorithms (35). In addition, prediction of neoantigen can contain inaccuracies due to multiple factors including the difficulty in modeling the processing and presentation of peptides by HLA(40). Improvement in the accuracy of neoantigen prediction and estimation TCR-neoantigen interaction strengths will be critical in predicting T cell clonal expansion driven by specific neoantigens using our model.

Our model is a minimalistic description of T cell clonal expansion initiated by neoantigens. T cell proliferation depends, in addition to the strength of TCR-pMHC interaction, on a variety of other factors such as co-stimulation of CD28 receptors, stimulation of inhibitory co-receptors such as PD1, CTLA4(41), and cytokine stimulation(42)- all of which are likely to be present in the TME. In the model these effects are captured by the estimated parameter *x_p_*. In addition, we made a simplifying assumption that each T cell spent the same amount of time in the TME until the samples were collected. Another simplifying assumption made was the T cells do not enter or leave the spatial region in the duration of interest T, the effect of different rates of T cell immigration and egress TME could be an interesting future direction. A more detailed prediction of T cell response such as expansion of a specific T cell in the TME will need to consider a more detailed modeling of the above processes.

## Materials and Methods

### 1. Development of the T cell clonal expansion model

*<u>Clonal expansion in the brain TME</u>:* A T cell#*i* waits a time interval τ before it encounters productive interactions from neoantigen-MHC molecules in a spatial region which leads to its proliferation in the time interval τ to T (**Figure 2E**). We consider the waiting time *τ* to be distributed in an exponentially increasing function with time in the inhibitory region, *Q*(𝜏) ∝ exp(𝑘𝜏), i.e., the longer the T cell waits the easier it becomes for it to experience a productive interaction to undergo proliferation. The waiting time distribution can be modeled by a Markov process (**Supplementary Text 1**). The stochastic proliferation process in the time interval *T-τ* is described by a Markov process where the probability for a T cell #*i* to proliferate in the time interval *t* to *t*+*dt* is given by *λr_i_dt*; *λ* is the proportionality constant relating the net strength of interaction *r_i_* with the propensity for T cell proliferation. Therefore, given T cell#i starts with a single cell initially (or 𝑚*_i_* = 1), the probability for a T cell #*i* to proliferate 𝑀*_i_* − 1 times in the time interval T-τ is ∝ exp(−𝜆*r* (𝑇 − 𝜏))(1 – exp(−𝜆*r_i_* (𝑇 − 𝜏)))*^M_i_^*^-1^. Since τ can range from 0 to *T*, the probability of finding a clone containing *M_i_* number of T cell #*i* in a spatial region is given by,

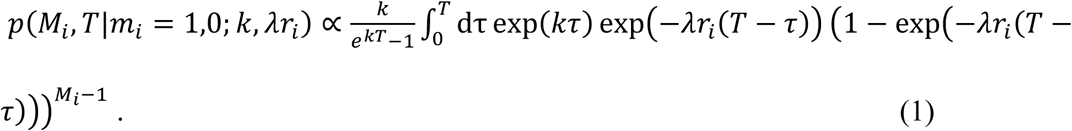

Further details regarding the derivation are provided in **Supplementary Text 1**.

The probabilities in Eq. (1) can be described in terms of dimensionless parameters {𝑥*_w_*, 𝑥*_p_*} defined as 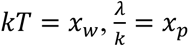.

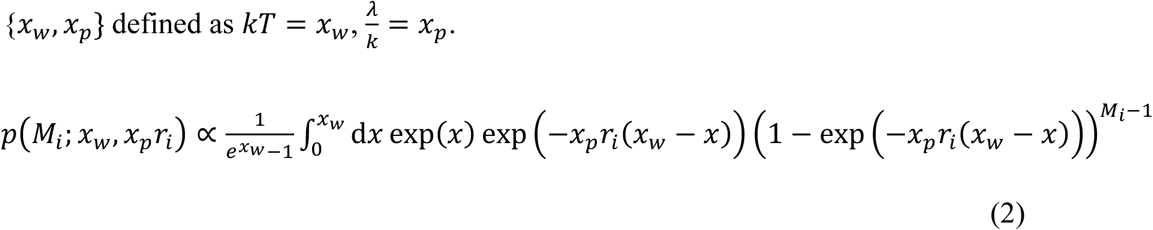

The probability in Eq. (2) can be shown to give rise to power law distributions i.e., 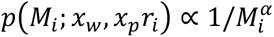, where *α* = 1 + 1/(*r_i_*𝑥*_p_*), under specific conditions such as when one type of neoantigen is present (**Supplementary Text 1**). We normalize the probability in Eq. (2) following 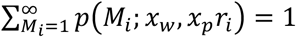 to create a likelihood function ℒ to estimate the model parameters given the clone sizes ({𝑀*_i_*}) and the proliferation rates {*r*_i_}. We divide {*r*_i_} by the largest value of {*r*_i_} across all the GBM and BrMET patients (or *r*_561_) to keep the values between 0 and 1. As a simplification, we assume that the formation of clones of different T cells are independent of each other. Therefore, ℒ is given by,

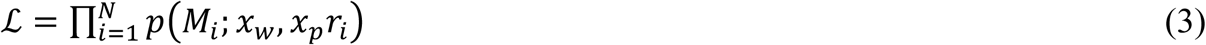

*<u>Clonal expansion in the periphery:</u>* Similar to the T cell proliferation model in the TME, we considered T cell proliferation to be stochastic where the propensity for proliferation for a T cell #*i* interacting with neoantigens{*u*} present in the vaccine is proportional to *r_i_* = ∑*_u_* 𝑠*_iu_* . The probability of finding a clone of size 𝑀_*i*_ post-vaccination for a T cell#*i*, given its size 𝑚_*i*_ pre-vaccine after a time duration *T* post-vaccination is given by,

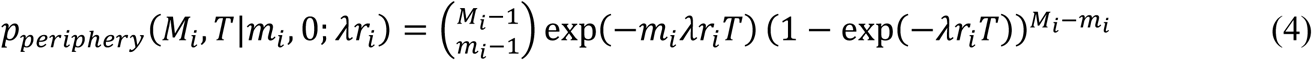

where 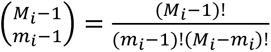 is the normalization constant.

As in the model for the TME, λ represents the proportionality constant relating the propensity for proliferation to *r*_*i*_. We estimated λ by maximizing the log-likelihood function ℒ*_periphery_*,

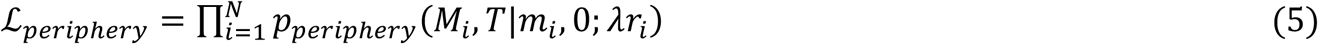

The likelihood functions in equations (2) and (5) can be described as

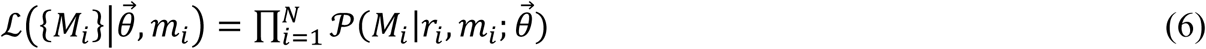

where parameter 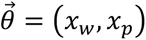 in the case of brain TME and 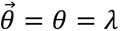 in the periphery. The best estimate of 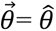 is given by the set of parameters which maximizes ℒ given the clone size data ({𝑀*_i_*}) and the estimated strengths of TCR-neoantigen-MHC or {*r_i_*}, i.e,

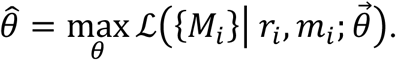

We obtain the *θ̂* by maximizing the log ℒ in Eq (6) or equivalently minimizing the negative of the log ℒ. We used “Nelder-Mead”(43) algorithm for such minimization. This algorithm is available is *scipy.optimize.minimize* package with a choice of an argument *method*.

### 2. Data acquisition and data pre-processing

*<u>TCRβ sequences</u>*: TCR𝛽 sequences and counts for GBM and BrMET patients (non-vaccinated) were obtained upon request from the authors of ref. (7). For the vaccine data case, the sequences and counts were obtained from Supplementary Table S5 in ref. (36). The neoantigen sequences were taken from Supplementary Table S3 provided in ref. (7) and Supplementary Table S4 in ref. (36). For TCR𝛽 sequences, if the counts were not available, they were omitted from the data. The WES data were downloaded from NCBI dbGaP for the dataset “General Research Use in Immunogenomics of Malignant Brain Tumors” associated with the ref. (7).

*<u>Estimation of</u>* <u>𝑠*_iu_ using PanPep and ERGO-II*</u>: We used zero-shot setting with all other default arguments for acquisition of 𝑠*<u>_iu_</u>* from *PanPep*. The command to run for this setting is *python PanPep.py --learning_setting zero-shot --input input.csv --output output.csv.* The input file contains two columns called ‘*Peptide*’ and ‘*CDR3*’. For a dataset (CDR3β and neoantigen sequences), each row would contain every possible pairing between the CDR3β sequences and neoantigen sequences. The output file would contain an additional third column ‘*Score*’ than that of input file which has the predicted binding score. If we index CDR3β sequences by 𝑖 and neoantigen sequences by 𝑢, then each row is a binding score 𝑠*<u>_iu_</u>*.

We used the same input files for *ERGO-II* prediction. *ERGO-II* comes with a choice of choosing a predictor based on which dataset it was trained on. The available options are ‘*VDJdb*’ and ‘*McPAS*’. We used both these training datasets for score predictions.

*Generation of clones from the best estimate θ̂*: For determining probabilities of occurrence for any T cell, we use probabilities in Eqs. (2) and (4) where the model parameters are set to the best estimate *θ̂*. We generated 1000 random configurations of T cell clone sizes for any spatial region using those probabilities in the following way. We chose a maximum clone size *M̃* = 10000(≫𝑀*_i_*) that might be generated in the random configuration and normalized the probability distributions in Eq. (2) and (4) assuming 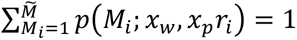 or 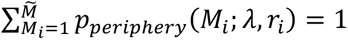. Then we randomly draw the T cells#𝑖 present in a spatial region using the normalized probability distributions. We generate 1000 such configurations. The clone size distributions are computed by evaluating the frequency *f_l_* of the clone size, *l,* given by *f_l_* = ∑*_i_* 𝛿*_l_*_,*mi*_ from the generated T cell configurations. Some of the clone sizes were generated only a few times (<10 times) in the 1000 configurations, we removed those clone sizes from the calculation of *f_l_.* For these selected clone sizes, we select a 95% of the probability mass at each selected clone sizes to construct a pointwise band. These bands are shown in probability distribution of clone sizes figures. For tutorial check out tutorial notebook in https://github.com/maheshworpaudel5001/TCR_proliferation_in_GBM.

## Data and code availability

The sample data and a tutorial are available at our GitHub page https://github.com/maheshworpaudel5001/TCR_proliferation_in_GBM.

## Acknowledgement

We thank Dr. Elaine Mardis for many stimulating discussions, insightful suggestions, and a critical review of the manuscript. We also thank Elizabeth A.R. Garfinkle, Katherine E. Miller, and Max Schaettler for their help with accessing the sequencing data used here. This work was supported by funding from the Nationwide Children’s Hospital.

## Author Contribution

Maheshwor Poudel: Conceptualization; investigation; methodology; software; validation; writing – original draft; writing – review and editing. William Stewart: Conceptualization; investigation; methodology; supervision; writing – original draft; writing – review and editing. Ciriyam Jayaprakash: Conceptualization; investigation; methodology; supervision; writing – original draft; writing – review and editing. Jayajit Das: Conceptualization; funding acquisition; investigation; methodology; project administration; supervision; writing – original draft; writing – review and editing.

