## Supplementary information for "Neoantigens and Stochastic Fluctuations Regulate T Cell Proliferation in Primary and Metastatic Malignant Brain Tumors"

PNAS

**This PDF file includes:**

Supporting text
Figures S1 to S7
Table S1

**Supplementary Text 1**

We model the proliferation of a T cell upon interacting with a pMHC as a simple birth or replication process, in which the number of clones, $M$ increases by one at a rate of $\lambda r$ for each clone.

$$M\,X \stackrel{M\lambda r}{\rightarrow} (M+1)X\,. \tag{1}$$

The net strength of the interaction is denoted by $r$ and $\lambda$ is the proportionality constant relating the strength to the rate. For notational clarity we have dropped the subscript $i$ used in the main text to refer to an individual T cell labelled by its unique TCR sequence $i$. This is an intrinsically stochastic process in which the randomness incorporates the inherent uncertainties in the various cell signaling and other biochemical processes. The stochastic dynamics of this birth process is described by the Master equation

$$\frac{dp_g(M,t)}{dt} = \lambda r\left[(M-1)p_g(M-1,t) - Mp_g(M,t)\right] \tag{2}$$

where $p_g(M,t)$ is the probability of obtaining $M$ clones at time in this simple process. If we assume that there is initially one clone, the probability of there being $M$ clones at time $t$, $p_g(M,T)$, is well-known to be given by the geometric distribution,

$$p_g(M,T) = e^{-\lambda rT}\left(1 - e^{-\lambda rT}\right)^{M-1}\,. \tag{3}$$

We assume that T-cells enter the TME and the tumor is resected after it has spent an average total duration of time $T$ in any spatial region. A T-cell has to wait a time interval $\tau$ to receive sufficient stimulatory signals in the immunosuppressive environment of the TME to start proliferating and therefore, in the model, it proliferates for a duration $T-\tau$. We assume that the waiting time $\tau$ that is determined by the local immunosuppressive environment to be a random variable.

So, if replication starts after a waiting time $\tau$, then the number of clones $M$ at $T$ obeys the distribution $p_g(T-\tau)$ defined above. This has to be averaged over the distribution of initial starting times $\tau$, $Q(\tau)$, and so, the number of clones $M$ at time $T$ obeys a doubly random process with a distribution given by

$$p(M,T) = \int_0^T d\tau\, Q(\tau)\, p_g(M,T-\tau)\,. \tag{4}$$

If $n$ positive encounters with antigens are needed for the T-cell to start proliferating then the waiting time $\tau$ is given by $\tau = \sum_{j=1}^{n} t_j$; if we assume that the $t_j$ are independently exponentially distributed, the probability distribution of $\tau$ for fixed $n$ is well-known and is the Gamma or Erlang distribution. The important feature of this distribution is that even for moderate values of $n$ the waiting time distribution is an increasing function of time for small $\tau$. For analytical tractability and computational feasibility of numerically fitting the data across different patients we choose a simple exponentially increasing form, $e^{k\tau}$ in the interval $[0,\, T]$. The normalized probability density is then given by

$$Q(\tau) \;=\; \frac{k}{e^{kT} - 1}\, e^{k\tau}\,. \tag{5}$$

Therefore, we have the result

$$p(M,T) \;=\; \frac{k}{e^{kT}\,-\,1} \int_0^T d\tau\, e^{k\tau}\, e^{-\lambda r\,(T-\tau)}\, \left[1 \,-\, e^{-\lambda r(T-\tau)}\right]^{M-1}\,. \tag{6}$$

We do a series substitutions that yields a more compact expression We first re-scale time by $k$, change the integration variable to $x \;\equiv\; k\tau$ and define $x_w \;\equiv\; kT$. Introducing the variable $x_p \;\equiv\; \frac{\lambda}{k}$ we find the following expression for $p(M,T)$.

$$p(M,T) \;=\; \frac{1}{e^{x_w}\,-\,1} \int_0^{x_w} dx\, e^x\, e^{-x_p r(x_w-x)}\, \left[1 \,-\, e^{-x_p r(x_w-x)}\right]^{M-1}\,. \tag{7}$$

This is Equation (2) in the main text.

To extract the power-law behavior it is convenient to define $e^{-x_p r(x_w-x)} \;\to\; u$. Making use of $\frac{du}{u} \;=\; x_p r dx$ and $e^{x-x_w} \;=\; u^{1/(x_p r)}$ and rearranging we have

$$p(M,T) \;=\; \frac{1}{1-e^{-x_w}}\, \frac{1}{x_p r} \int_{e^{-x_p r x_w}}^{1} du\, u^{1/(x_p r)}\, (1 \,-\, u)^{M-1}\,. \tag{8}$$

Provided $e^{-x_p r x_w}$ is small the lower limit can be set equal to zero and the integral can be expressed in terms of Gamma functions:

$$\int_0^1 du\, u^{1/(x_p r)}\, (1 \,-\, u)^{M-1} \;=\; \frac{\Gamma\left(1 \,+\, \frac{1}{x_p r}\right)\, \Gamma(M)}{\Gamma\left(M + 1 \,+\, \frac{1}{x_p r}\right)}\,. \tag{9}$$

Now, we can extract the $M$ dependence since

$$\ln\Gamma(z) \;\sim\; \left(z \,-\, \frac{1}{2}\right) \ln z \,-\, z \,+\, \frac{1}{2}\, \ln(2\pi) \,+\, O(1/z) \tag{10}$$

for large $M$ and we find $p(M,\, T) \;\sim\; M^{-1-\frac{1}{x_p r}}$. This exponent is larger in magnitude than 1 and is independent of $T$.

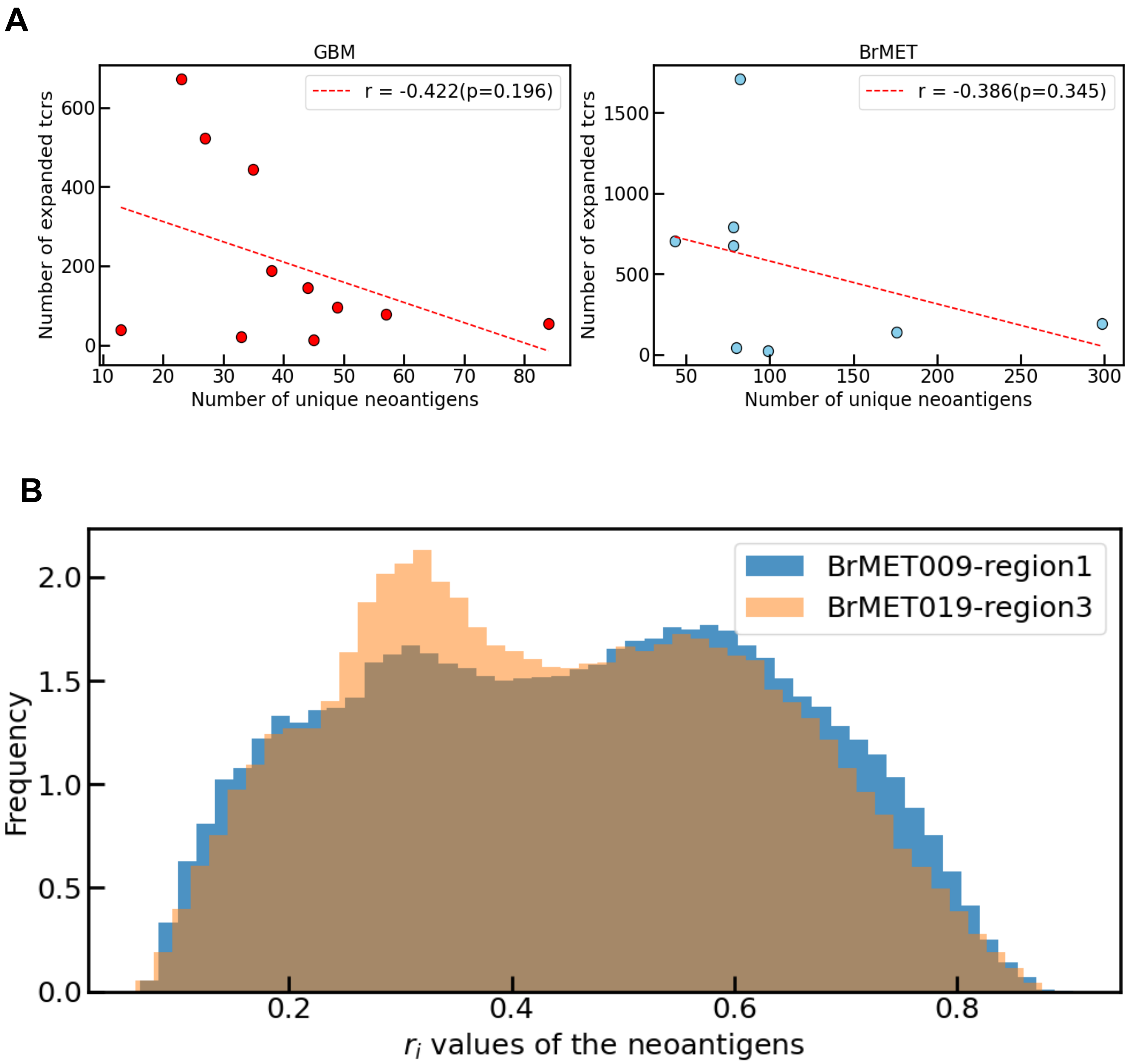


**Figure S1. (A)** Shows correlation between the number of distinct TCR clones that expanded (with counts >=5) with the number of distinct neoantigens in GBM (left panel) and BrMET (right panel) patients as reported by Schaettler et al.. The correlations are not statistically significant. This is in contrast to positive correlation of the number of distinct T cell clones and the number of non-synonymous mutations reported for NSCLC tumors in ref. (18). (B) Shows distributions of the estimated propensity $\{r_i\}$ values in two tumor regions of BrMET patient where region 1 in BrMET009 and region 2 in BrMET019 contain 72 and 289 numbers of distinct neoantigens, respectively. The distributions largely overlap though one region (region 1) contains 4 times more neoantigens but 22 times lesser number of distinct TCRs than the other region (region 2). The lack of correlation in (A) could be due to the differences in the strength of interactions between neoantigens and TCRs in different tumor regions.

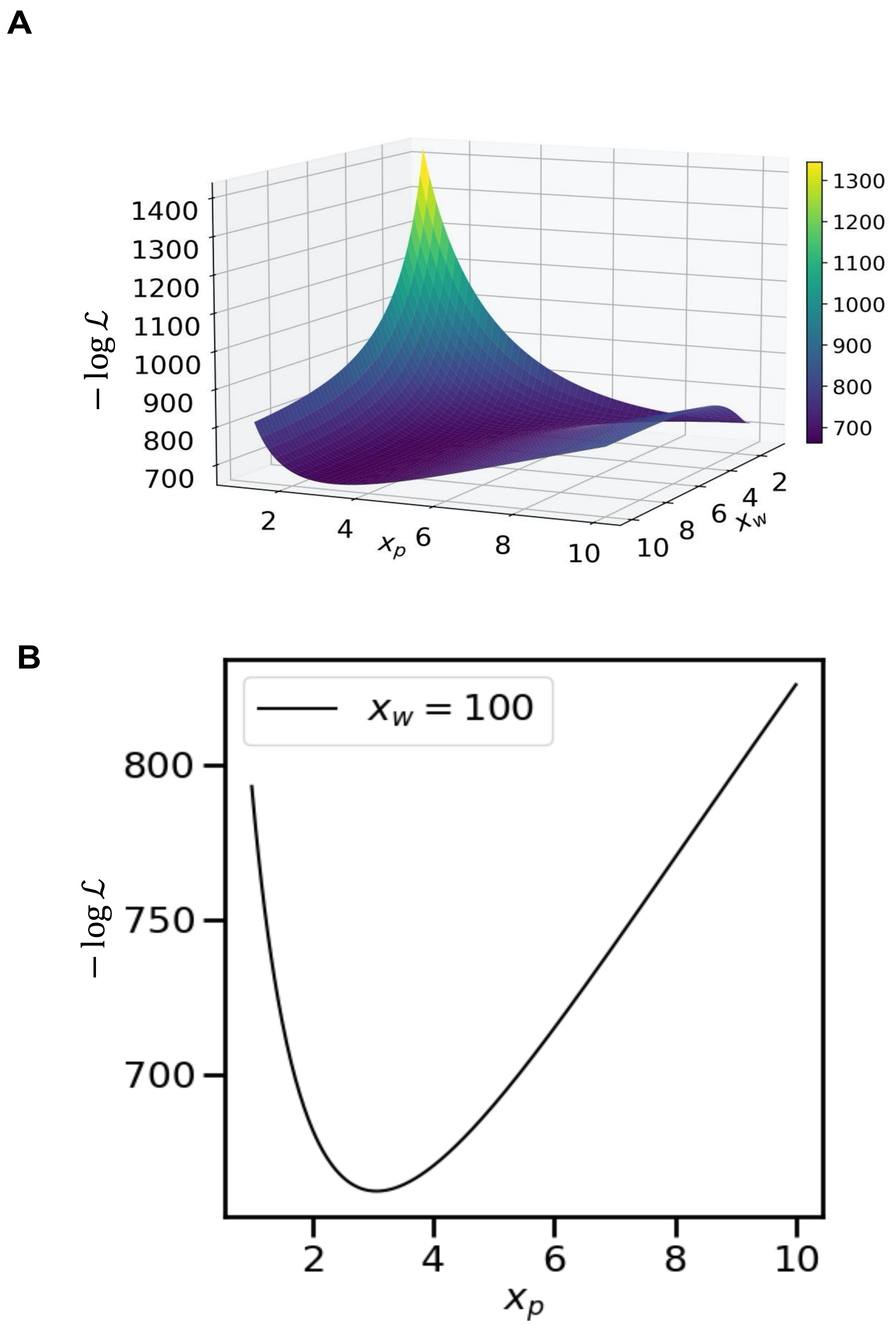


**Figure S2. Variation of the Likelihood $L$ with the model parameters $x_w$ and $x_p$.**
**(A)** Shows the variation of $-\log \mathcal{L}$ with the parameters $x_w$ and $x_p$ for the T cell clone data in region 1 of patient BrMET008. $-\log \mathcal{L}$ changes more with $x_p$ than $x_w$. In our estimation of $x_p$ we fixed $x_w$ to 100. The parameter $x_p$ for all the data set is provided in Supplementary table1. **(B)** The variation of $-\log \mathcal{L}$ with $x_p$ when $x_w$ is fixed to 100.

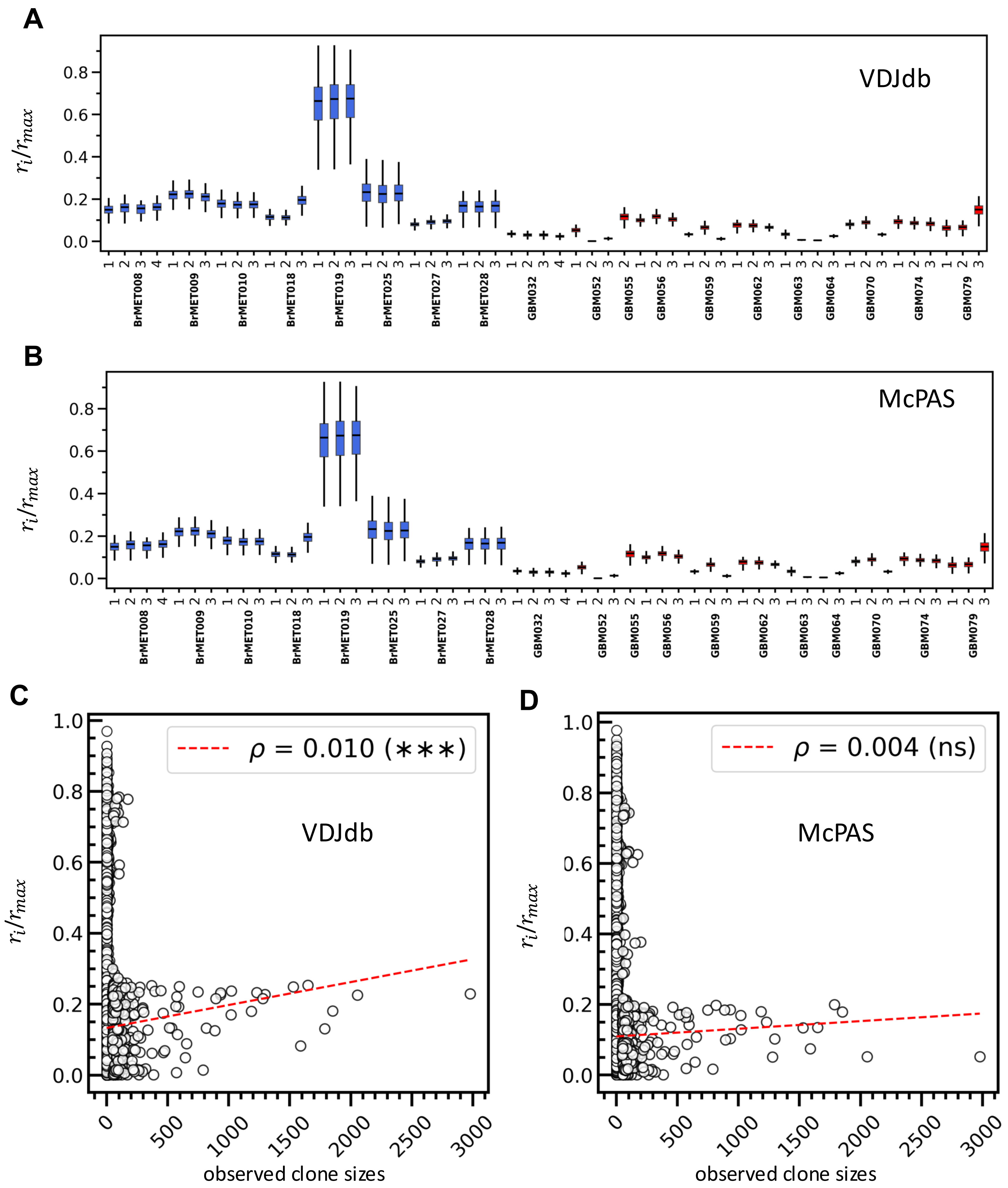


**Figure S3.** Variation of T cell proliferation rates estimated by ERGO-II in different spatial regions. (A and B) Box plot of scaled T cell proliferation rate, $r_i$, estimated using ERGO-II trained on (A) VDJdb data set and trained on (B) McPAS data set for all BrMET patients (in blue) and GBM patients (in red). For an individual patient, the values do not show appreciable differences within a spatial region. The results are qualitatively similar to that obtained using PanPep (Figure 2B, main text). (C and D) Scatter plot showing variation of $r_i$ estimated using ERGO-II and the size of the clone for T cell#i. ERGO-II trained on VDJdb dataset (C) and on McPAS dataset (D) are shown. The correlation is negligible ($\leq 0.01$) similar to the results obtained for the estimations performed by PanPep (Figure 2D, main text). Regression line shown in red. Significance determined from Pearson R correlation test. Although, the test signifies that the correlation between these two quantities is significant, however the correlation is negligible

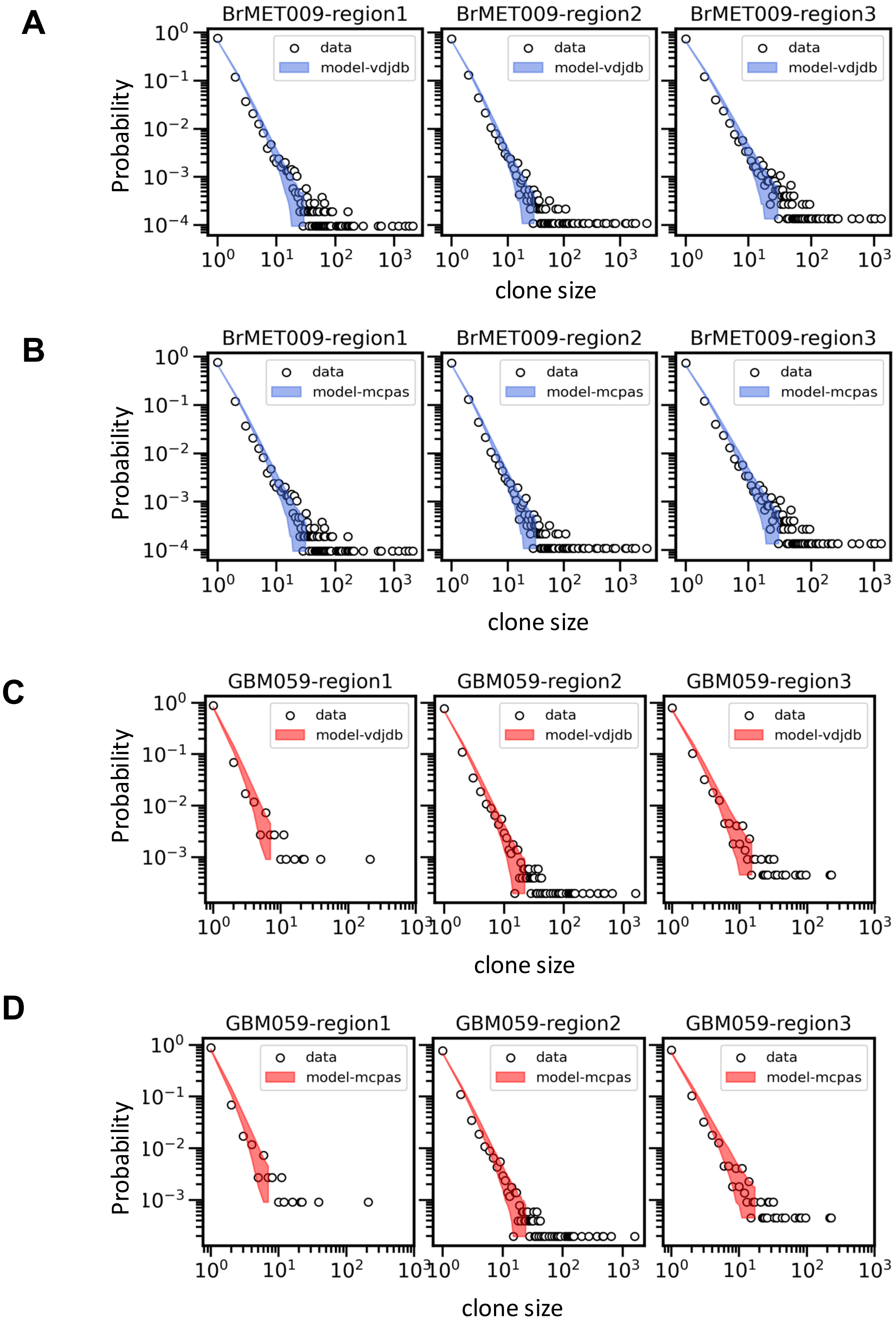


**Figure S4. Model prediction for clone size distributions when T cell proliferation rates are estimated by *ERGO-II*.** Model prediction of the clone sizes for BrMET patient BrMET009 (A-B) and GBM patient GBM059 (C-D). *ERGO-II* trained on *VDJdb* (A, C) and *McPAS* (B, D) were used to calculate $\{r_i\}$ for the estimation of model parameters. The quality of the prediction is similar to that obtained for the case when the $\{r_i\}$ were estimated using *PanPep* (**Figure 3D**, main text).

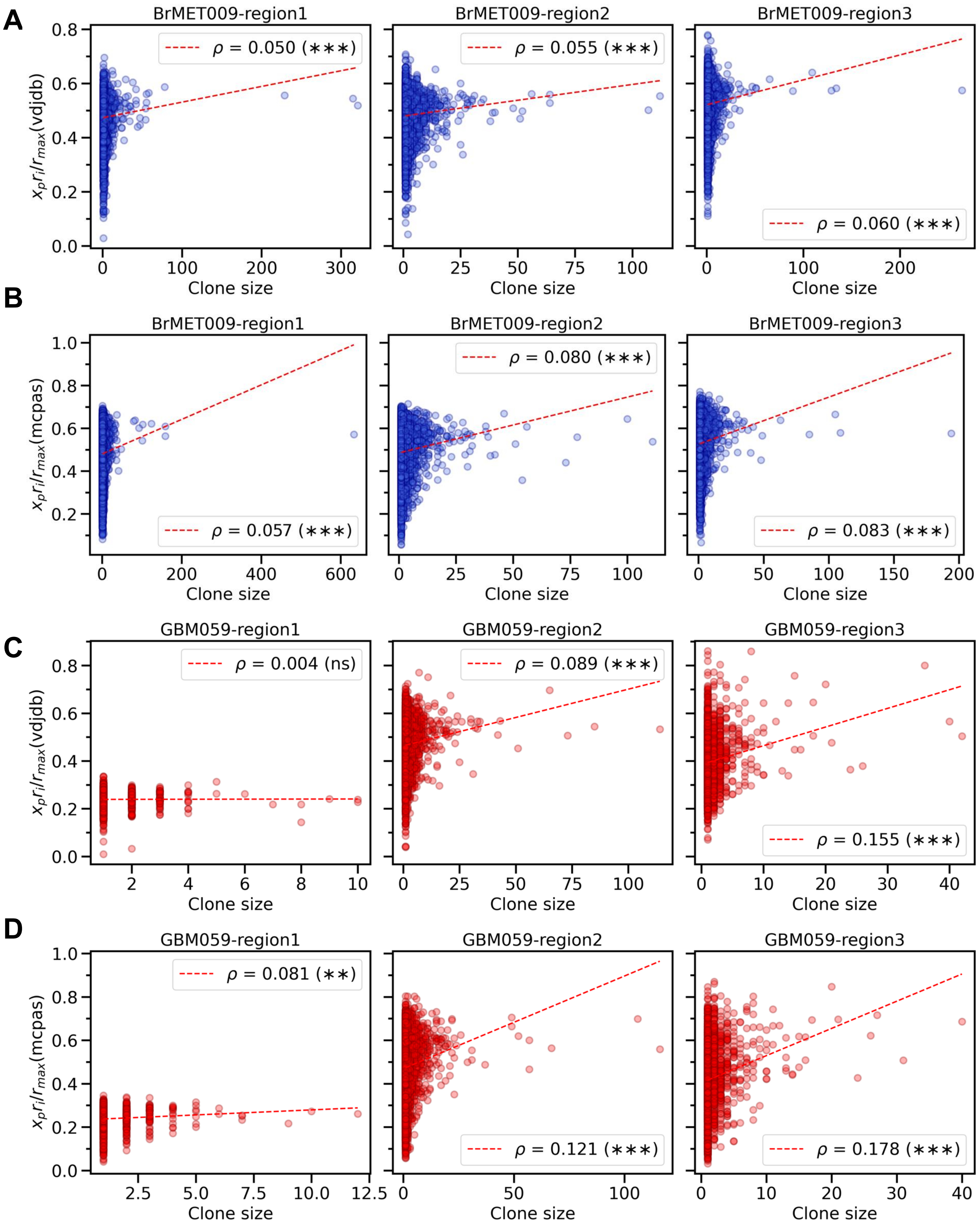


**Figure S5. Correlation between the T cell proliferation rate estimated using *ERGO-II* and the T cell clone size.** Shows correlation between the scaled rate of proliferation $r_i$, evaluated for *ERGO-II* trained on trained on (A) *VDJdb* data set and trained on (B) *McPAS* data set for for BrMET009. Correlations between the same variables for GBM059 are shown in for *ERGO-II* trained on (C) *VDJdb* or (D) *McPAS*. The results are qualitatively the same to that obtained using *PanPep* (**Figure 3F**, main text).

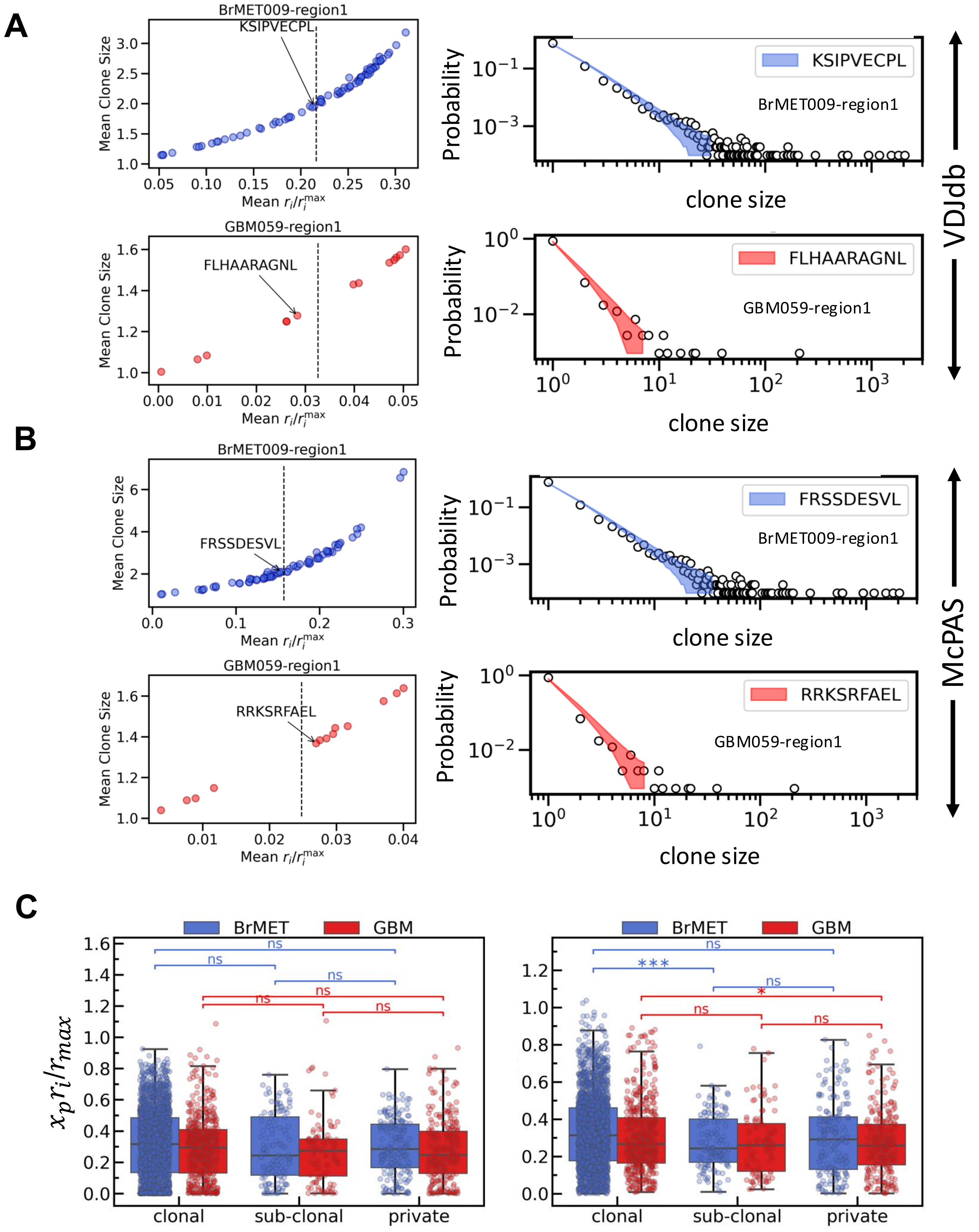


**Figure S6. Prediction of the contribution of different neoantigens to generate T cell expansion when the T cell proliferation rate is estimated using *ERGO-II*.**

Contribution of individual neoantigens is producing T cell expansion is estimated following the scheme shown in Figure 4 (main text). **(A-B) (left panels)** Shows a wide range of mean T cell clone sizes produced by different neoantigens. (**right panels**) The neoantigens that produce mean T cell clone sizes similar to that of the observed mean T cell clone sizes also generate similar clone size distributions compared to the observed T cell clone size distributions. *ERGO-II* was trained on (A) *VDJdb* or (B) *McPAS*. The above results are qualitatively similar to that shown in **Figure 5B** in the main text where the results are obtained using *PanPep*. However, the rank ordering of neoantigens, and the specific neoantigens that generate mean T cell clone sizes similar to that of the observed mean T cell clone sizes are different for *ERGO-II* compared to that for *PanPep*. This shows the differences in *ERGO-II* and *PanPep* in estimating TCR-pMHC interactions which arise from the inherent inaccuracies in the estimations of the interactions can make predictions for specific neoantigens challenging. (C) Box plots of mean propensity for T cell proliferation for neoantigens estimated using *ERGO-II* (*VDJdb* left) and (*McPAS* right) for BrMET and GBM groups across regions differentiated as three categories clonal, sub-clonal and private neoantigens based on their presence in the region sequenced. The clonal neoantigens show higher mean propensity than the other neoantigens irrespective of the type of tumor. Significance determined by two-sided Welch's t-test. The results are qualitatively similar to that obtained using *PanPep* shown in **Figure 4C** in main text.

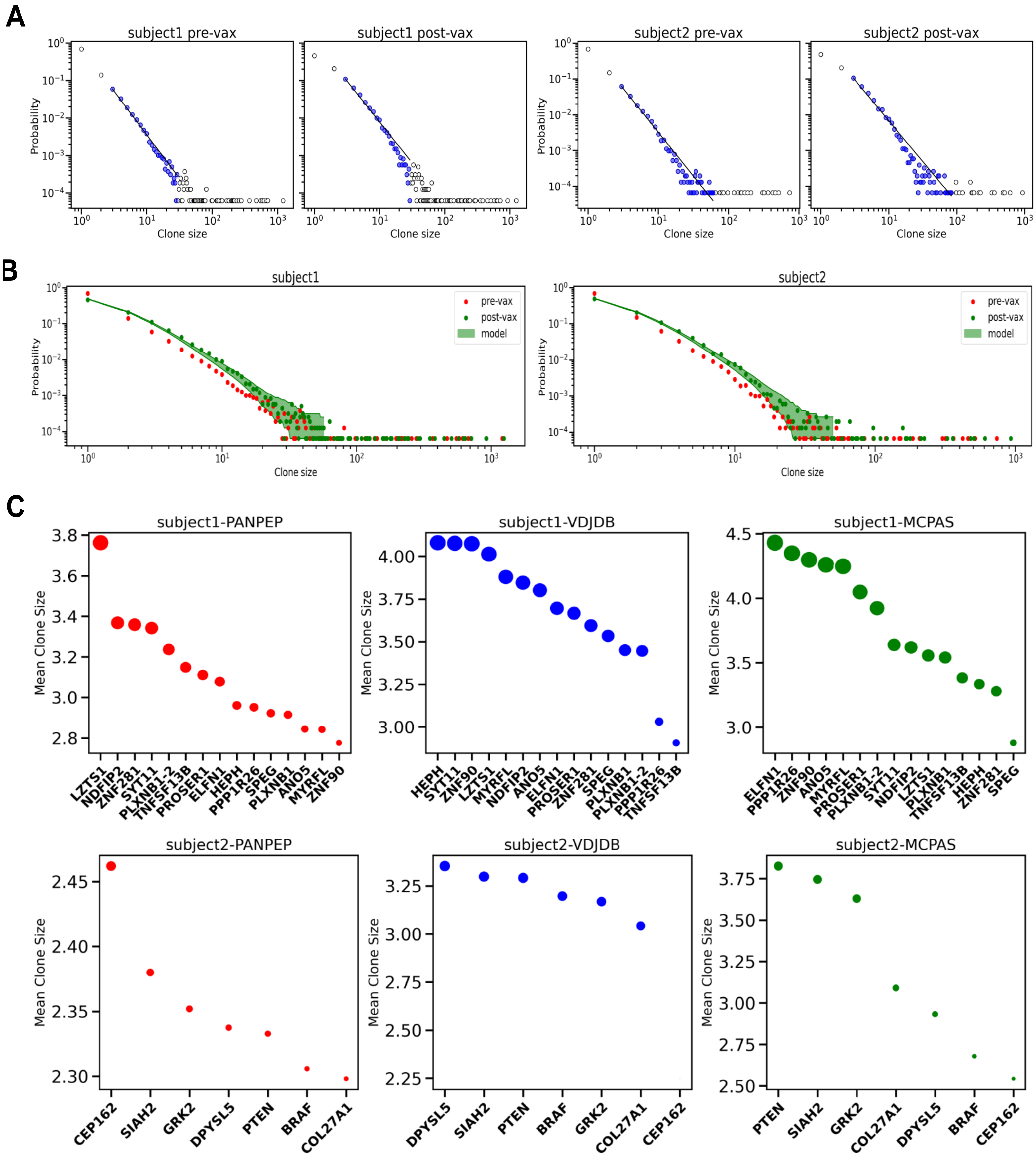


**Figure S7. Clonal expansion of T cells in response to neoantigen vaccines in glioblastoma patients, subject#2 and subject#3.**

**(A)** Power law fit to probability distribution of observed clone sizes (solid black line to blue dots) in subject 1 and 2 pre- and post-vaccination. **(B)** Comparison of model predictions of T cell clone sizes in subject 1 and 2. **(C)** Rank ordering (ascending) of synthetic long peptides used in neoantigen vaccines for subject 1 and 2 according to the model predicted mean clone size when mixture of neoantigens is replaced by a neoantigen one at a time. The size of the dots represents the magnitude of mean propensity for the respective peptide. The order depends on the relative estimates of the TCR-pMHC strengths estimated by the bioinformatic tools (*PanPep* and *ERGO-II*) we used.

| **Table S1.** Parameter values obtained from minimizing negative log likelihood | | | |
|---|---|---|---|
| Patient | lambda_panpep | lambda_vdjdb | lambda_mcpas |
| subject1 | 0.0087 | 0.006 | 0.006 |
| subject2 | 0.0081 | 0.015 | 0.023 |
| subject3 | 0.0046 | 0.005 | 0.008 |
